# Estimation of Left Ventricular Mechanical Activation Times from Motion-Corrected Cardiac 4DCT Images

**DOI:** 10.1101/2022.04.12.488106

**Authors:** Ashish Manohar, James Yang, Jed D. Pack, Elliot R. McVeigh

**Affiliations:** Department of Mechanical and Aerospace Engineering, University of California San Diego, La Jolla, California, USA; Department of Bioengineering, University of California San Diego, La Jolla, California, USA; Radiation Systems Lab, GE Global Research, Niskayuna, New York, USA; Department of Radiology, University of California San Diego, La Jolla, California, USA; Department of Medicine, Cardiovascular Division, University of California San Diego, La Jolla, California, USA

**Author notes:** **Corresponding Author:** Elliot McVeigh, PhD, 9452 Medical Center Drive, La Jolla, CA 92037.

**Keywords:** cardiac CT, motion correction, LV mechanical activation, cardiac resynchronization therapy

## Abstract

**Background:** Mechanical activation times of the left ventricle (LV) are strongly correlated with patient response to cardiac resynchronization therapy (CRT); however, there remains an unmet clinical need for a simple, robust imaging modality that yields accurate and reproducible estimates of LV mechanical activation times. Modern four-dimensional computed tomography (4DCT) imaging systems with effective motion correction could fulfill this need.

**Purpose:** To demonstrate the clinical utility of a newly developed cardiac motion correction algorithm called ResyncCT and to investigate its effect on estimating mechanical activation times of the LV.

**Methods:** Twenty-four subjects with full cardiac volume, full cardiac cycle 4DCT images were retrospectively analyzed in this study. The reconstructed 4DCT images exhibited typical motion artifact characteristics that were dependent on the direction of LV wall motion with respect to the gantry position at each cardiac phase; these artifacts rotated synchronously with the gantry. The motion corrupted images, referred to as the uncorrected images, were processed with a novel cardiac motion correction algorithm called ResyncCT to yield motion corrected images (referred to as the ResyncCT images). Regional shortening (RS_CT_) of the LV was calculated over a 72-segment model for both the uncorrected and the ResyncCT images; each segment contained RS_CT_ vs time data. From these RS_CT_ vs time curves, LV mechanical activation was estimated as the time point at which the RS_CT_ vs time curve shortened by 10% of its full dynamic range during systolic contraction; we refer to this time point as the time to onset of shortening (TOS). A shift in TOS > |35| ms has a significant effect on reclassifying patients based on their probabilities of responding to CRT; we investigated the effect of ResyncCT on the measured values of TOS over the LV.

**Results:** ResyncCT had a pronounced effect on the TOS estimates; ResyncCT could potentially reclassify 23/24 (96%) subjects as either responders or non-responders to CRT. In 16/24 (67%) subjects, the differences in TOS that were sufficient to bring about a reclassification of CRT response were at least as large in surface area of the LV as one entire American Heart Association segment.

**Conclusions:** We demonstrated the clinical utility of ResyncCT in estimating LV mechanical activation times as ‘times to onset of shortening’ in 24 human subjects. ResyncCT has a pronounced effect on the estimation of LV mechanical activation times; the differences in activation times between the ResyncCT and uncorrected images are heterogenous and subject specific. The effect of ResyncCT could potentially reclassify 96% of the subjects used in this study based on their probabilities of responding to CRT. The results reported in this study highlight the potential utility of ResyncCT in estimating timing of mechanical events of interest of the LV for CRT planning.

## 1. Introduction

Cardiac resynchronization therapy (CRT) is an effective treatment for patients in heart failure and with left ventricular (LV) dyssynchrony [1]; however, 30-50% of patients selected for CRT do not respond to the treatment [2]. Current guidelines for CRT patient selection include echocardiography-derived left ventricular ejection fractions (LVEF) ≤ 35%, New York Heart Association (NYHA) functional classes II-IV, and QRS durations > 120 ms [3]. Significant effort has been focused on reducing this non-responder rate through more effective patient selection with the use of non-invasive imaging, primarily with echocardiography, but with limited success thus far [4], [5].

Cardiac magnetic resonance (CMR) has been shown to be an excellent modality for the accurate and reproducible estimation of timing of mechanical events of the LV [6]–[8]. CMR-derived estimates of LV mechanical activation have also been shown to be strongly correlated with CRT response [9]–[11]; thus, establishing LV mechanical activation as an important parameter for predicting CRT response. However, the complexity and limited availability of highly skilled CMR centers has hindered its routine clinical use. Additionally, 28% of patients under consideration for CRT already have existing right ventricular (RV) pacing systems in place, serving as a contraindication for CMR imaging in many of these patients [3].

Four-dimensional x-ray computed tomography (4DCT) can yield high-resolution 3D volumetric images across the cardiac cycle with low radiation dose [12]. The images can also be acquired using routine FDA approved scanning protocols available from all vendors which do not require specially trained personnel. Previous studies have successfully used dual-source 4DCT for LV dyssynchrony assessment and CRT planning [13]–[15]. Dual-source scanners offer higher temporal resolution; however, due to their limited z-axis detector coverage, they can suffer from step-artifacts that arise during helical acquisition of the superior-inferior extent of the heart over multiple irregular heartbeats. We previously reported that out of 147 subjects recruited for a retrospective CRT study, 37 subjects (25%) had severe step-artifacts preventing the precise estimation of LV mechanics [16]. With the advances made in 4DCT motion correction technology [17], wide detector scanners can now yield high temporal resolution images, together with the advantages of single-heartbeat and single-table position acquisitions [18]; the ‘false dyssynchrony’ artifact [19], [20] can also be significantly reduced [21]. These favorable features of 4DCT are of particular value in the accurate and precise estimation of LV mechanical activation times, highlighting the potential utility of wide detector 4DCT systems in guiding CRT planning.

In addition to the CMR-derived estimates of LV mechanical activation, we have also previously shown that the ‘time to onset of shortening’ (TOS) at the proposed implantation site of the LV left lead is a very important feature in predicting CRT response [16]. Thus, the objective of this study was to assess the effect of a newly developed cardiac CT motion correction algorithm called ResyncCT on the computed TOS estimates in a clinical cohort of 25 subjects. ResyncCT has previously been validated using a set of controlled phantom experiments [21]; this study presents new results which focus on the clinical applicability and the effect of ResyncCT on routinely acquired retrospective cardiac 4DCT images in human subjects.

## 2. Methods

### 2.1 Subjects

Twenty-five consecutive subjects that underwent 4DCT exams between January 2018 and July 2021 for the evaluation and planning of transcatheter aortic valve replacement (TAVR) and that were clinically diagnosed as having “normal LV function” were included in this study. The scans were acquired and read by radiologists at the University of California San Diego (UCSD). Neither subject enrolment, image acquisition, image reconstruction, nor clinical diagnosis was modified for the purpose of this particular study; the subjects were scanned and images diagnosed as per routine clinical protocols established at UCSD for pre-TAVR evaluation. The de-identified images of these subjects were retrospectively used in this study in accordance with an IRB approved protocol.

### 2.2 4DCT Imaging and Left Ventricle Segmentation

The 25 subjects were scanned with a 256-detector row scanner (Revolution CT, General Electric Healthcare, Chicago, IL) under an established clinical imaging protocol for TAVR assessment at UCSD. The protocol included retrospective ECG-gated single-heartbeat full cardiac cycle imaging with no x-ray tube current modulation applied. The 256-detector row scanner has a z-axis coverage of 16 cm, permitting full heart volume imaging from a single table position. The gantry revolution time was 280 ms. The scans were acquired with tube voltages of 80 kVp (n = 2), 100 kVP (n = 21), and 120 kVp (n = 2) and tube currents of 340 mA (n = 2), 440 mA (n = 2), 600 mA (n = 3), 660 mA (n = 2), and 720 mA (n = 16). The images were reconstructed at 70 ms intervals (90 degrees of rotation for a gantry revolution time of 280 ms) using the ‘Standard’ reconstruction kernel of the Revolution CT into 512 x 512 x 256 voxels. The in-plane pixel spacings were in the range of [0.31, 0.53] mm with a slice thickness of 0.625 mm for all images.

The reconstructed 4DCT images of each subject exhibited motion artifacts that are dependent on the direction of motion of the endocardial walls with respect to gantry position; these artifacts rotate synchronously with the orientation of the gantry, giving a false impression of dyssynchronous contraction [19]. The motion uncorrected images are referred to hereafter as the *uncorrected* images. These gantry position induced artifacts make it very difficult to accurately measure the mechanical activation time of regions of the LV because of the uncertainty of endocardial wall position in the images. The uncorrected images were processed with a motion correction algorithm called ResyncCT [21] to yield motion corrected images. ResyncCT is a newly developed cardiac CT motion correction algorithm that leverages the power of conjugate pairs of partial angle reconstruction images for motion estimation and motion compensation. The motion corrected images are referred to hereafter as the *ResyncCT* images. Figure 1 shows an axial CT slice in an example subject with and without motion correction demonstrating the clarification of endocardial wall position in the corrected image.

**Fig. 1.**
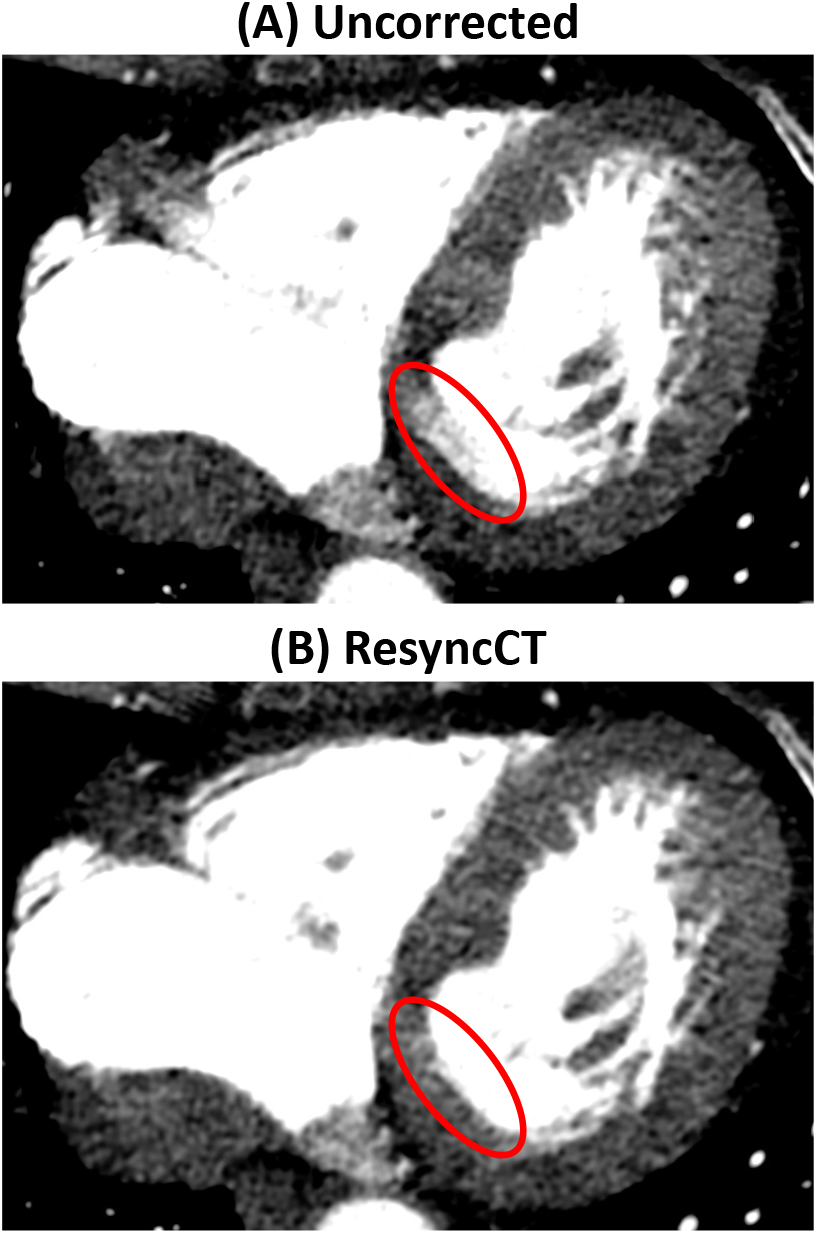
Reconstructed axial CT slice in an example subject showing motion artifact. (**A**) Uncorrected CT image showing pronounced “double-wall” motion artifact highlighted by the red ellipse. (**B**) ResyncCT image showing significant reduction of the “double-wall” artifact, and clarification of the position of the endocardial wall.

For each subject, the LV blood volume was segmented from each time frame of the uncorrected and the ResyncCT image series. The segmentation procedure has been previously described in detail [22]. Briefly, the segmentation threshold was determined using Otsu’s method [23] and the active contour region growing module of ITK-SNAP v3.8.0 [24] was used to segment the LV blood volume. The volume of the LV for each time frame was computed by summing the segmented 3D voxels. Meshes delineating the LV endocardium were then extracted from the segmented LV volumes.

### 2.3 Endocardial Regional Shortening

Endocardial regional shortening (RS_CT_) [25]–[27] was estimated by registering the end diastolic LV mesh to each subsequent mesh derived from the other time frames of the cardiac cycle using a point set registration technique [28] and was computed according to the following equation:

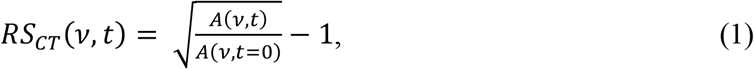

where *A*(*y,t*) is the area of an endocardial patch *v* at time *t* of the cardiac cycle; time *t* = 0 corresponds to end diastole. The patch *v* is defined on the end-diastolic mesh -it is tracked as a “material point” through the subsequent time frames. The average patch size for a normal human LV was 2.6 ± 0.5 mm^2^ yielding a high-resolution LV map of regional endocardial shortening.

The high resolution endocardial surface map of RS_CT_ was divided into 72 endocardial segments: 18 in the circumferential direction (1 segment for every 20°) for each of 4 slices from apex to base along the LV long axis [22]; all RS_CT_ values within each segment were averaged to yield a single RS_CT_ value for that segment. This 72-segment model yields a higher spatial sampling of endocardial regional shortening than the traditional AHA 16-segment model [29]; the higher sampling is in agreement with the superior resolution of LV features visible on the 4DCT images [30], [31]. Each segment corresponds to approximately 2 cm^2^ of endocardial surface for a normal sized LV. Two 72 segment models were constructed for each subject in this study: 1) derived from the original uncorrected CT images and 2) derived from the ResyncCT images.

### 2.4 Measuring the mechanical activation time: Time to Onset of Shortening (TOS)

The TOS was estimated for all 72 segments of the LV for both the uncorrected and the ResyncCT images. The TOS for a particular segment was calculated as the time at which the RS_CT_ vs time curve for that segment shortened by 10% of its full dynamic range during systolic contraction. We previously investigated estimating mechanical activation times of LV wall motion with 4DCT using controlled phantom experiments [21]. The main results of that study highlighted that the estimates of activation times are more accurate and precise when measured during the constant motion profile of the LV during systolic contraction, vs. the time of the “pre-stretch peak” which was previously used as the mechanical activation time in tagged MR studies of dyssynchrony [32], [33]. Figure 2 illustrates the estimation of the TOS for an example RS_CT_ vs time curve from one LV segment.

**Fig. 2.**
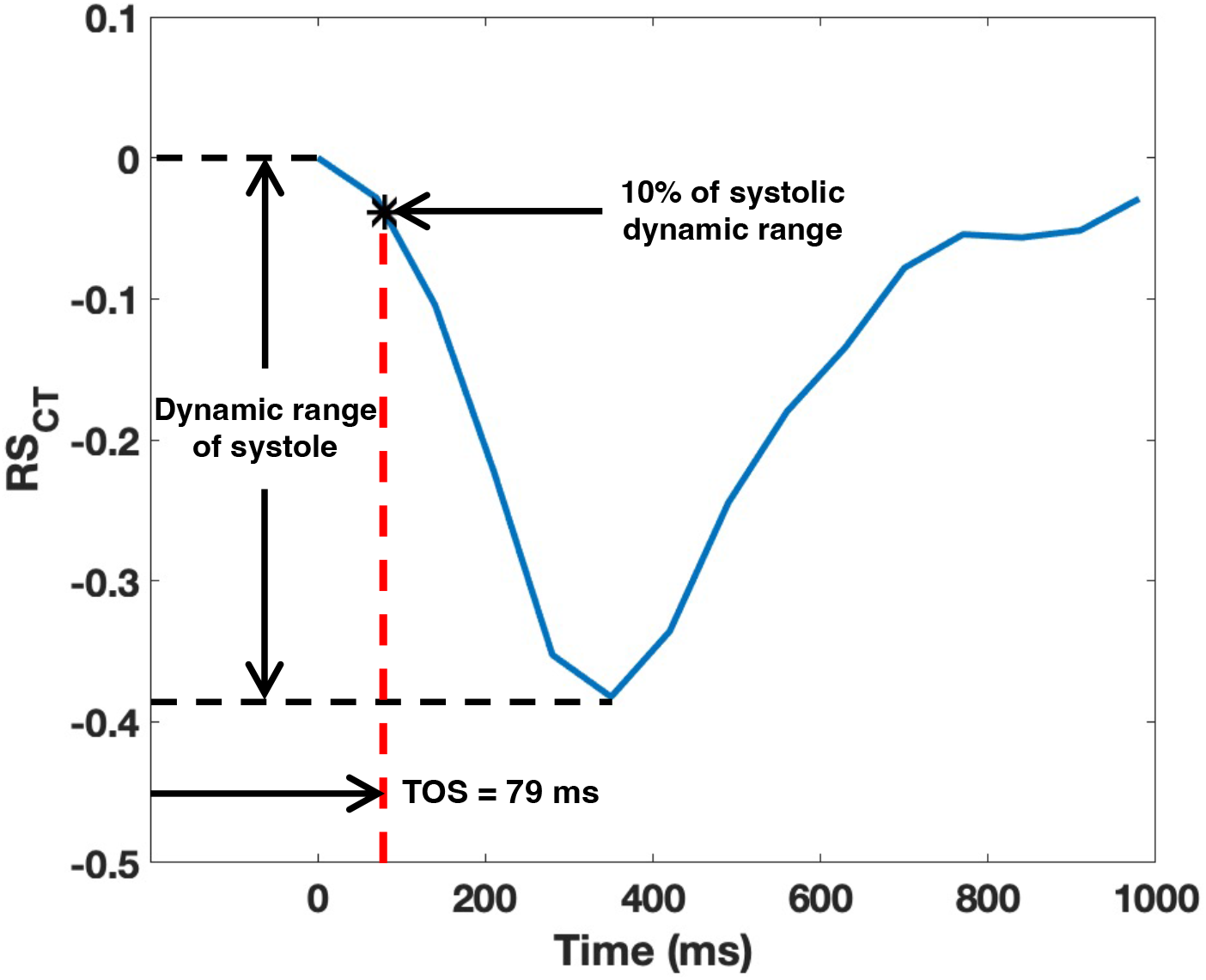
Estimation of the time to onset of shortening (TOS) for an example RS_CT_ vs time curve. The TOS has been shown to be the most accurate metric for estimating mechanical activation time. RS_CT_: endocardial regional shortening.

### 2.5 Significance of Time to Onset of Shortening in Predicting CRT Response

The TOS has previously been shown to be an excellent metric to quantify LV dyssynchrony [6]-[8] and is also strongly correlated with CRT response [9]—[11]. From our previous study, it was determined that a shift in TOS (|ΔTOS|) of 35 ms will cause a significant shift in the classification of patients between low, mid, and high probability of CRT response [16]. In this study, the effect of ResyncCT was evaluated by observing the number of LV segments (as % LV surface area) that experienced a change in TOS estimates by greater than |35| ms between the uncorrected and the ResyncCT images for each subject.

## 3. Results

### 3.1 Subject Characteristics

Out of the 25 subjects used in this study, one had to be excluded due to significant calcification around the mitral valve, preventing the precise definition of the mitral valve plane. Ten of the 24 subjects were female (42%), and the average age of the entire cohort was 79 ± 9 years (median: 77 years; interquartile range: 14 years). The mean CT-derived LVEF was 69.8 ± 8% and 69.5 ± 8% for the uncorrected and the ResyncCT images, respectively (p = 0.88). Table 1 lists the subject characteristics.

**Table 1.** Subject characteristics.

|  |  |  |
| --- | --- | --- |
| **N** | 24 |  |
| **Age (years)** | $79 \pm 9$ | |
| **Female, n (%)** | 10 (42) |  |
| **Heartrate (bpm)** | $67 \pm 13$ | |
| **4DCT-derived EF (%)** | **Uncorrected** | $69.8 \pm 8$ |
| | **ResyncCT** | $69.5 \pm 8$ |
| **4DCT-derived volumes (mL)** | **EDV** | $147 \pm 57$ |
| | **ESV** | $48 \pm 32$ |
| **DLP (mGy\*cm)** | $509 \pm 252$ | |

bpm: beats per minute; EF: ejection fraction; EDV: end diastolic volume; ESV: end systolic 220 volume; DLP: dose length product

### 3.2 False Dyssynchrony

Figure 3 highlights the pronounced effect of motion correction with ResyncCT on the position of the LV walls across the cardiac cycle. Figure 3A shows axial slices of the ResyncCT-derived images in an example subject for five consecutive time frames of the cardiac cycle. Figures 3B and 3C show difference images between the consecutive time frames of the uncorrected and the ResyncCT images, respectively. In the uncorrected images, for a particular time frame, walls moving perpendicular to the x-ray beam direction at the time of sampling are updated while the walls moving in a direction parallel to the x-rays are not updated; therefore, pairs of walls are updated only every other time frame in a series of 4DCT images reconstructed at 90° intervals of gantry rotation (70 ms for a gantry rotation time of 280 ms). Figure 3C highlights the continuous motion field recovered uniformly over all regions of the LV by ResyncCT.

**Fig. 3.**
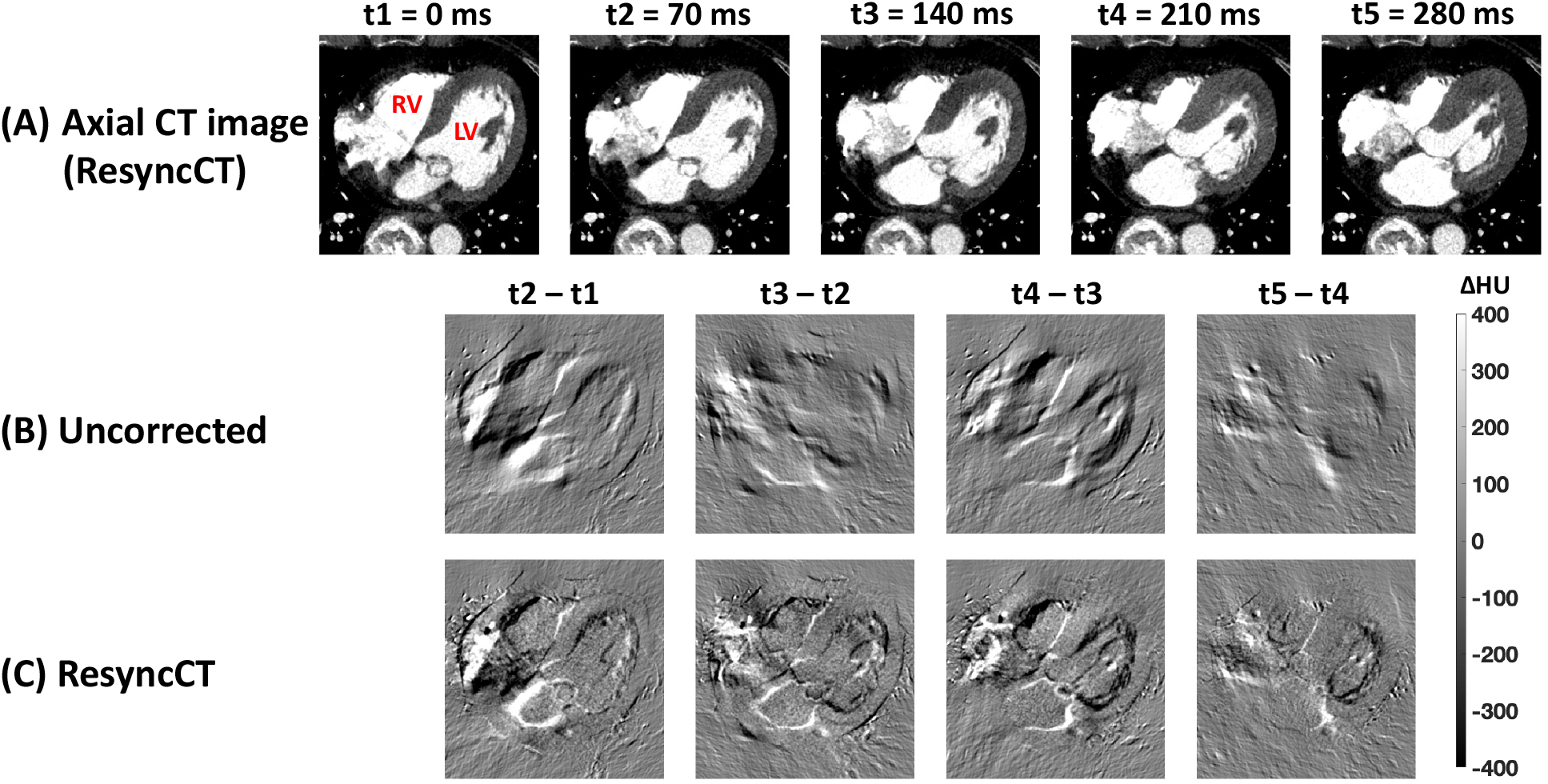
False dyssynchrony and recovery of motion field with ResyncCT in an example subject. (**A**) Axial slices of the ResyncCT-derived images in an example subject for five consecutive time frames across the cardiac cycle. (**B-C**) Difference images between the consecutive time frames of (**B**) the uncorrected images (not shown) and (**C**) the ResyncCT images shown in (A). Pairs of myocardial walls are updated every other time frame in the uncorrected images whereas ResyncCT recovers a continuous motion over all LV regions at each time frame. Note the clarity of the edges, and position of the right coronary artery in the ResyncCT processed images. LV: left ventricle; RV: right ventricle.

### 3.3 Effect of ResyncCT on Accuracy of Mechanical Activation Time: Estimates of Time to Onset of Shortening

Figures 4A–4B show two example subjects and the effect of ResyncCT on their TOS estimates. For each subject, the first row shows the high-resolution bullseye maps of TOS, the second row shows mid-LV short-axis slices for three time frames of the cardiac cycle, and the third row shows difference images of the short-axis slices between the first three time frames. The figures on the left are derived from the uncorrected images and the figures on the right are derived from the ResyncCT images. Figure 4A shows a subject whose uncorrected images-derived TOS map has higher values on the anteroseptal wall as compared to those obtained from the ResyncCT images. Due to gantry-induced artifacts, the motion field between time frames t3 and t2 is lost in the uncorrected images as seen in the difference image on the bottom left; however, ResyncCT recovers this motion field. Figure 4B shows a subject with higher TOS values derived from the uncorrected images on the inferolateral wall; again, as seen in the difference image between time frames t3 and t2 of the uncorrected images on the bottom left, the inferolateral wall of the LV is not well localized due to gantry-induced artifacts.

**Fig. 4.**
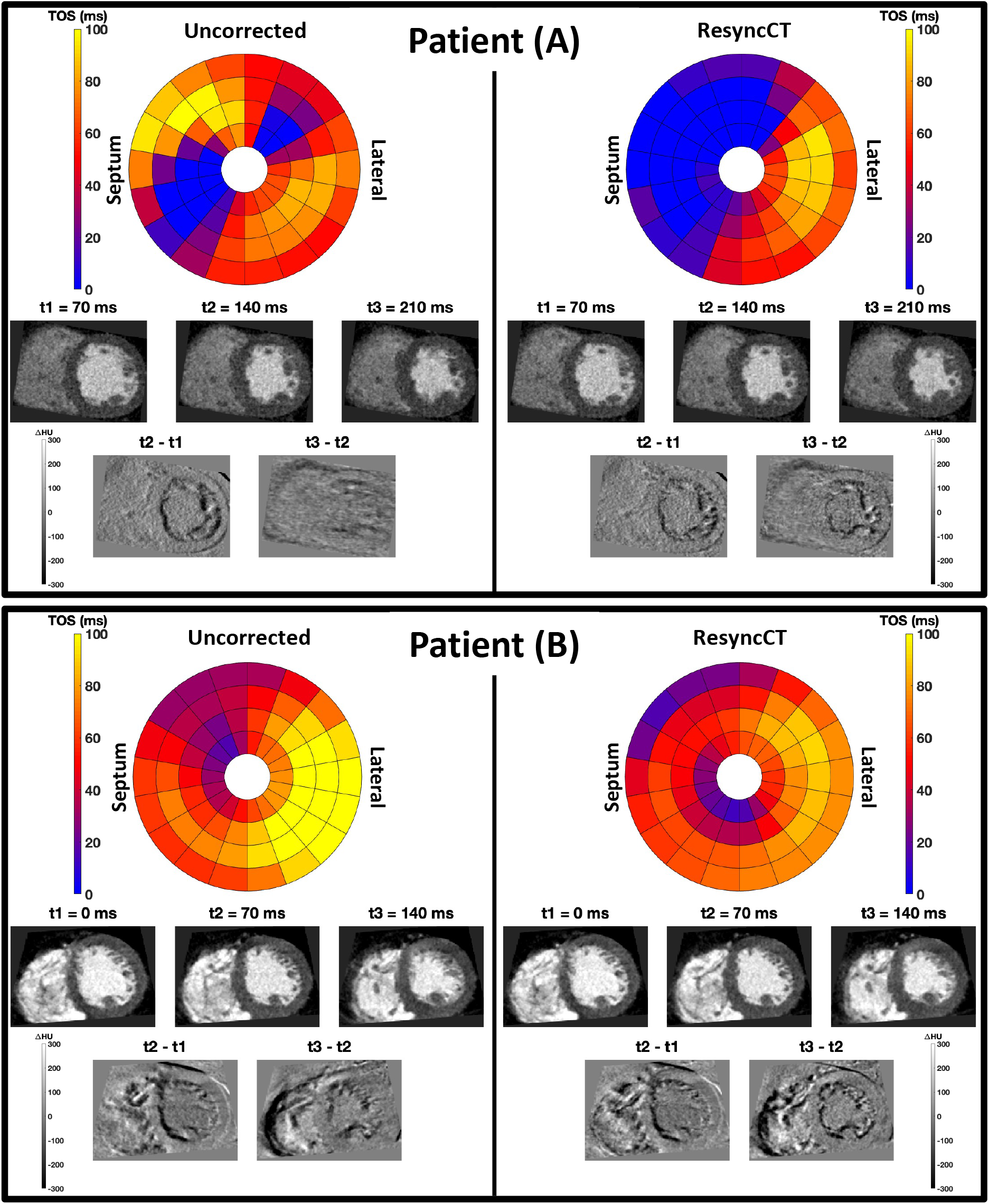
Subject specific effects of ResyncCT on the estimation of TOS in two example subjects (A) and (B). The left column gives results from uncorrected images, and the right column are results in the same subject using ResyncCT corrected images. In each column, the first row shows bullseye maps of TOS, the second row shows mid-LV short-axis slices for three time frames of the cardiac cycle, and the last row shows difference images between the first three time frames of the mid-LV short-axis images. TOS: time to onset of shortening. The bullseye plot views the LV from the apex to base.

Only one subject had no part of their endocardial surface differ in TOS estimates ≥ |35| ms; thus, in 23/24 (96%) subjects, ResyncCT had a significant impact on the TOS map and could potentially reclassify these subjects based on their probability of responding to CRT. Figure 5A shows bullseye maps of the difference in TOS estimates between the uncorrected and the ResyncCT images over all 72 endocardial segments for all 24 subjects; virtually all of the patients have substantial changes to the TOS values. Figure 5A also highlights the heterogenous subject-specific differences in the TOS estimates between the uncorrected and the ResyncCT images; the degree of motion correction achieved depends on the direction of motion of the endocardial walls with respect to the position of the gantry. Figure 5B shows the distribution of the percentage of the LV endocardial surface that differed in TOS estimates ≥ |35| ms between the uncorrected and the ResyncCT images. Sixteen subjects (67%) had 7 or more endocardial segments differ in TOS estimates ≥ |35| ms; the 7 segment threshold approximately corresponds to one of the standard AHA 16 segments [29], or 10% of the LV endocardial surface (14 cm^2^).

**Fig. 5.**
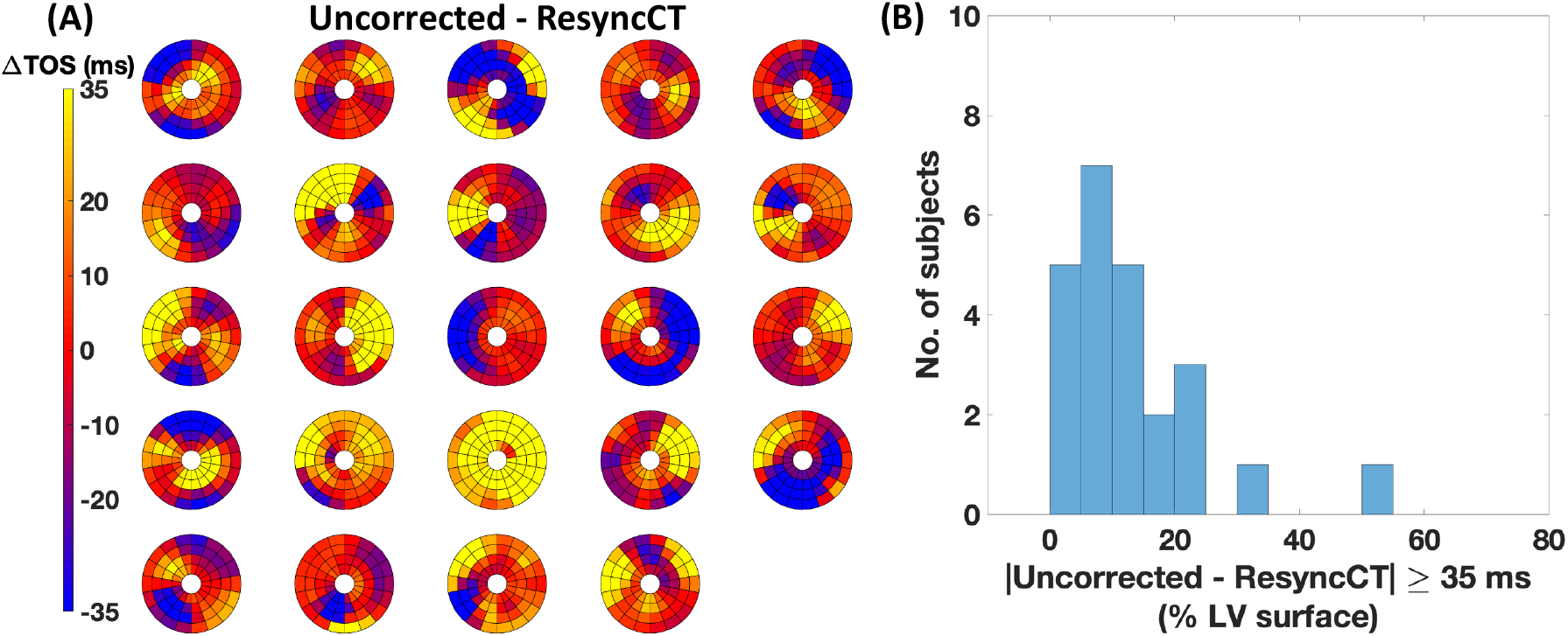
Effect of ResyncCT on TOS estimates over the LV. (**A**) Bullseye maps of the difference in TOS estimates across the entire LV between the uncorrected and the ResyncCT images for all 24 subjects. (**B**) Histogram showing the number of subjects vs the percentage of LV endocardial surface that differs in TOS estimates ≥ |35| ms between the uncorrected and the ResyncCT images. LV: left ventricle; TOS: time to onset of shortening.

## 4. Discussion

In this study, we evaluated the ability of a newly developed cardiac CT motion correction algorithm called ResyncCT to remove heart wall motion artifacts on routinely acquired clinical 4DCT images. The main findings of this study reveal the clinical utility of ResyncCT in minimizing gantry-induced false dyssynchrony artifacts in cardiac 4DCT images; this is especially important when estimating the mechanical activation time of the myocardium as ‘times to onset of shortening’ from cardiac 4DCT images of subjects under consideration for CRT. In 23/24 subjects in this study, ResyncCT could have a significant effect in potentially reclassifying the subjects based on their probability of responding to CRT [16]. The TOS is the most prominent feature in determining the probability of CRT response; a change in TOS of greater than 35 ms could significantly change the estimated probability of a subject responding to CRT; thus, the accurate and precise estimation of TOS from 4DCT is essential.

We have previously evaluated the effect of ResyncCT under controlled phantom experiments [21], the results from which highlight the excellent accuracy and precision of ResyncCT-derived TOS estimates of LV wall motion. The work reported here expands that validation of ResyncCT to clinically acquired human 4DCT studies. Another important finding in our previous phantom studies was that the accuracy and precision of the TOS estimates were higher when measured on the constant motion profile of LV wall motion during systole vs. measuring the peak of pre-stretch; previous studies with tagged MRI used the peak of the strain curves to characterize mechanical activation delays [7], [33]. For this reason, the TOS in our study was defined as the time of the cardiac cycle where the RS_CT_ vs time curve reached 10% of its dynamic range during systolic contraction.

This study also highlights the heterogenous effect of “gantry artifact” on the TOS estimates of the 24 subjects; LV orientation, heart rate, LV systolic motion profile (velocity and acceleration), and gantry position with respect to LV wall motion during systole are some of the factors that make the corrections of ResyncCT subject specific. It is impossible to predict the interaction of a patient’s heartrate with the gantry orientation, so ResyncCT is needed in case there are profound motion artifacts.

The TOS estimates are derived from RS_CT_ vs time data which has previously been shown to have very low inter- and intra-observer variability The analysis pipeline used in this study to estimate high-resolution maps of TOS over the entire LV was fully automated, except for the definition of the mitral valve and LVOT planes. The inter- and intra-observer variability of the segmentation procedure, and in turn the variability of the RS_CT_ estimates has previously been shown to be very low [22], making the estimation of TOS from RS_CT_ vs time data to be highly reproducible. Additionally, with advances in deep learning cardiac image segmentation [34], [35], we expect the LV blood pool segmentation to be fully automated in the near future. This will ensure that results are exceptionally reproducible.

While dual-source CT scanners yield images with higher temporal resolution, their limited z-axis coverage of the detectors (~4 cm) poses difficulties for full-heart volume imaging of patients with irregular heartbeats and/or arrythmias. The patients must be scanned in helical or “step and shoot” mode over multiple beats which may then yield step-artifacts, often rendering the images unanalyzable for the estimation of timing of mechanical events of the LV. Wider detector scanners (256- or 320-detector rows) with effective motion estimation and motion compensation technologies may be better suited for this application.

### 4.1 Limitations

This study was successful in highlighting the need for subject-specific application of ResyncCT on the estimates of TOS in a cohort of 24 subjects. While the results reveal that ResyncCT can potentially reclassify 96% of the subjects based on their probability of responding to CRT, the study lacks a ground-truth value of TOS to validate the accuracy of these estimates. However, our previous study validating the accuracy and precision of the ResyncCT-derived TOS estimates [21] with ground-truth phantom experiments provides evidence supporting the validity of the estimates reported in this clinical cohort. The clinical utility of the results reported in this study, together with our previous validation using phantom experiments, motivate future studies to perform a direct comparison of the ResyncCT-derived TOS estimates with either cine CMR or CMR tagging.

All 24 subjects were scanned using the same CT scanner under the same clinical imaging protocol established at the center. The imaging protocol did not include any tube-current modulation across the cardiac cycle. While this was beneficial in estimating TOS values that weren’t subject to image quality differences across the cardiac cycle, future studies should investigate the effect of tube-current modulation and ultra-low-dose scan protocols on the computed TOS values.

## 5. Conclusions

A novel cardiac CT motion correction algorithm called ResyncCT was shown to dramatically reduce motion artifacts on the endocardial wall in clinically acquired 4DCT studies. Mechanical activation times evaluated by the time to onset of regional shortening, a parameter shown to be a strong predictor of CRT response, were significantly changed after motion correction with ResyncCT.

## Acknowledgements

Research reported in this publication was supported by NHLBI of the National Institutes of Health under award number R01 HL144678. The content is solely the responsibility of the authors and does not necessarily represent the official views of the National Institutes of Health.

This work was done during the term of an Award from the American Heart Association (AHA 20PRE35210261).

## References

[1] F. W. Prinzen, K. Vernooy, and A. Auricchio, “Cardiac Resynchronization Therapy,” Circulation, vol. 128, no. 22, pp. 2407–2418, Nov. 2013, doi: 10.1161/CIRCULATIONAHA.112.000112.

[2] K. Vernooy, C. J. M. van Deursen, M. Strik, and F. W. Prinzen, “Strategies to improve cardiac resynchronization therapy,” Nat. Rev. Cardiol., vol. 11, no. 8, pp. 481–493, Aug. 2014, doi: 10.1038/nrcardio.2014.67.

[3] J.-C. Daubert et al., “2012 EHRA/HRS expert consensus statement on cardiac resynchronization therapy in heart failure: implant and follow-up recommendations and management: A registered branch of the European Society of Cardiology (ESC), and the Heart Rhythm Society; and in col,” Europace, vol. 14, no. 9, pp. 1236–1286, Sep. 2012, doi: 10.1093/europace/eus222.

[4] V. Delgado and J. J. Bax, “Assessment of Systolic Dyssynchrony for Cardiac Resynchronization Therapy Is Clinically Useful,” Circulation, vol. 123, no. 6, pp. 640–655, Feb. 2011, doi: 10.1161/CIRCULATIONAHA.110.954404.

[5] R. K. Sung and E. Foster, “Assessment of Systolic Dyssynchrony for Cardiac Resynchronization Therapy Is Not Clinically Useful,” Circulation, vol. 123, no. 6, pp. 656–662, Feb. 2011, doi: 10.1161/CIRCULATIONAHA.110.954420.

[6] E. R. McVeigh, F. W. Prinzen, B. T. Wyman, J. E. Tsitlik, H. R. Halperin, and W. C. Hunter, “Imaging asynchronous mechanical activation of the paced heart with tagged MRI,” Magn. Reson. Med., vol. 39, no. 4, pp. 507–513, Apr. 1998, doi: 10.1002/mrm.1910390402.

[7] B. T. Wyman, W. C. Hunter, F. W. Prinzen, and E. R. McVeigh, “Mapping propagation of mechanical activation in the paced heart with MRI tagging,” Am. J. Physiol. Circ. Physiol., vol. 276, no. 3, pp. H881–H891, Mar. 1999, doi: 10.1152/ajpheart.1999.276.3.H881.

[8] B. T. Wyman, W. C. Hunter, F. W. Prinzen, O. P. Faris, and E. R. McVeigh, “Effects of single-and biventricular pacing on temporal and spatial dynamics of ventricular contraction,” Am. J. Physiol. Circ. Physiol., vol. 282, no. 1, pp. H372–H379, Jan. 2002, doi: 10.1152/ajpheart.2002.282.1.H372.

[9] D. A. Auger et al., “Imaging left-ventricular mechanical activation in heart failure patients using cine DENSE MRI: Validation and implications for cardiac resynchronization therapy,” J. Magn. Reson. Imaging, vol. 46, no. 3, pp. 887–896, Sep. 2017, doi: 10.1002/jmri.25613.

[10] K. C. Bilchick et al., “Impact of Mechanical Activation, Scar, and Electrical Timing on Cardiac Resynchronization Therapy Response and Clinical Outcomes,” J. Am. Coll. Cardiol., vol. 63, no. 16, pp. 1657–1666, Apr. 2014, doi: 10.1016/j.jacc.2014.02.533.

[11] R. J. Taylor, F. Umar, J. R. Panting, B. Stegemann, and F. Leyva, “Left ventricular lead position, mechanical activation, and myocardial scar in relation to left ventricular reverse remodeling and clinical outcomes after cardiac resynchronization therapy: A feature-tracking and contrast-enhanced cardiovascular magnetic r,” Hear. Rhythm, vol. 13, no. 2, pp. 481–489, Feb. 2016, doi: 10.1016/j.hrthm.2015.10.024.

[12] M. Y. Chen, S. M. Shanbhag, and A. E. Arai, “Submillisievert Median Radiation Dose for Coronary Angiography with a Second-Generation 320–Detector Row CT Scanner in 107 Consecutive Patients,” Radiology, vol. 267, no. 1, pp. 76–85, Apr. 2013, doi: 10.1148/radiol.13122621.

[13] J. Gould et al., “Feasibility of intraprocedural integration of cardiac CT to guide left ventricular lead implantation for CRT upgrades,” J. Cardiovasc. Electrophysiol., vol. 32, no. 3, pp. 802–812, Mar. 2021, doi: 10.1111/jce.14896.

[14] Q. A. Truong et al., “Utility of dual-source computed tomography in cardiac resynchronization therapy—DIRECT study,” Hear. Rhythm, vol. 15, no. 8, pp. 1206–1213, Aug. 2018, doi: 10.1016/j.hrthm.2018.03.020.

[15] D. B. Fyenbo et al., “Transmural Myocardial Scar Assessed by Cardiac Computed Tomography,” J. Comput. Assist. Tomogr., vol. 43, no. 2, pp. 312–316, 2019, doi: 10.1097/RCT.0000000000000824.

[16] A. Manohar et al., “Prediction of CRT Response Using a Lead Placement Score Derived from 4DCT,” medRxiv, p. 2022.03.23.22272846, Mar. 2022, doi: 10.1101/2022.03.23.22272846.

[17] A. Z. Kyme and R. R. Fulton, “Motion estimation and correction in SPECT, PET and CT,” Phys. Med. Biol., vol. 66, no. 18, p. 18TR02, Sep. 2021, doi: 10.1088/1361-6560/ac093b.

[18] S. Kim, Y. Chang, and J. B. Ra, “Cardiac motion correction based on partial angle reconstructed images in x-ray CT,” Med. Phys., vol. 42, no. 5, pp. 2560–2571, Apr. 2015, doi: 10.1118/1.4918580.

[19] M. Kidoh et al., “False dyssynchrony: problem with image-based cardiac functional analysis using x-ray computed tomography,” in Medical Imaging 2017: Physics of Medical Imaging, Mar. 2017, no. 10132, p. 101321U, doi: 10.1117/12.2250257.

[20] F. Contijoch, J. W. Stayman, and E. R. McVeigh, “The impact of small motion on the visualization of coronary vessels and lesions in cardiac CT: A simulation study,” Med. Phys., vol. 44, no. 7, pp. 3512–3524, Jul. 2017, doi: 10.1002/mp.12295.

[21] A. Manohar, J. D. Pack, A. J. Schluchter, and E. R. McVeigh, “Four-dimensional computed tomography of the left ventricle, Part II: Estimation of mechanical activation times,” Med. Phys., Mar. 2022, doi: 10.1002/mp.15550.

[22] G. M. Colvert et al., “Novel 4DCT Method to Measure Regional Left Ventricular Endocardial Shortening Before and After Transcatheter Mitral Valve Implantation,” Struct. Hear., vol. 5, no. 4, pp. 410–419, Jul. 2021, doi: 10.1080/24748706.2021.1934617.

[23] N. Otsu, “A Threshold Selection Method from Gray-Level Histograms,” IEEE Trans. Syst. Man. Cybern., vol. 9, no. 1, pp. 62–66, Jan. 1979, doi: 10.1109/TSMC.1979.4310076.

[24] P. A. Yushkevich et al., “User-guided 3D active contour segmentation of anatomical structures: Significantly improved efficiency and reliability,” Neuroimage, vol. 31, no. 3, pp. 1116–1128, Jul. 2006, doi: 10.1016/j.neuroimage.2006.01.015.

[25] E. R. McVeigh et al., “Regional myocardial strain measurements from 4DCT in patients with normal LV function,” J. Cardiovasc. Comput. Tomogr., vol. 12, no. 5, pp. 372–378, Sep. 2018, doi: 10.1016/j.jcct.2018.05.002.

[26] A. Manohar, G. M. Colvert, A. Schluchter, F. Contijoch, and E. R. McVeigh, “Anthropomorphic left ventricular mesh phantom: a framework to investigate the accuracy of SQUEEZ using Coherent Point Drift for the detection of regional wall motion abnormalities,” J. Med. Imaging, vol. 6, no. 04, p. 1, Dec. 2019, doi: 10.1117/1.JMI.6.4.045001.

[27] F. J. Contijoch, D. W. Groves, Z. Chen, M. Y. Chen, and E. R. McVeigh, “A novel method for evaluating regional RV function in the adult congenital heart with low-dose CT and SQUEEZ processing,” Int. J. Cardiol., vol. 249, pp. 461–466, Dec. 2017, doi: 10.1016/j.ijcard.2017.08.040.

[28] A. Myronenko and Xubo Song, “Point Set Registration: Coherent Point Drift,” IEEE Trans. Pattern Anal. Mach. Intell., vol. 32, no. 12, pp. 2262–2275, Dec. 2010, doi: 10.1109/TPAMI.2010.46.

[29] M. D. Cerqueira et al., “Standardized Myocardial Segmentation and Nomenclature for Tomographic Imaging of the Heart,” Circulation, vol. 105, no. 4, pp. 539–542, Jan. 2002, doi: 10.1161/hc0402.102975.

[30] A. Manohar et al., “Regional dynamics of fractal dimension of the left ventricular endocardium from cine computed tomography images,” J. Med. Imaging, vol. 6, no. 04, p. 1, Nov. 2019, doi: 10.1117/1.JMI.6.4.046002.

[31] J. P. Cruz-Bastida et al., “Hi-Res scan mode in clinical MDCT systems: Experimental assessment of spatial resolution performance,” Med. Phys., vol. 43, no. 5, pp. 2399–2409, Apr. 2016, doi: 10.1118/1.4946816.

[32] F. W. Prinzen, W. C. Hunter, B. T. Wyman, and E. R. McVeigh, “Mapping of regional myocardial strain and work during ventricular pacing: experimental study using magnetic resonance imaging tagging,” J. Am. Coll. Cardiol., vol. 33, no. 6, pp. 1735–1742, May 1999, doi: 10.1016/S0735-1097(99)00068-6.

[33] O. P. Faris et al., “Novel Technique for Cardiac Electromechanical Mapping with Magnetic Resonance Imaging Tagging and an Epicardial Electrode Sock,” Ann. Biomed. Eng., vol. 31, no. 4, pp. 430–440, Apr. 2003, doi: 10.1114/1.1560618.

[34] C. Chen et al., “Deep Learning for Cardiac Image Segmentation: A Review,” Front. Cardiovasc. Med., vol. 7, no. March, Mar. 2020, doi: 10.3389/fcvm.2020.00025.

[35] D. M. Vigneault, F. Contijoch, C. P. Bridge, K. Lowe, C. Jan, and E. R. McVeigh, “M-SiSSR: Regional Endocardial Function Using Multilabel Simultaneous Subdivision Surface Registration,” in Functional Imaging and Modeling of the Heart, 2021, pp. 242–252, doi: 10.1007/978-3-030-78710-3_24.

